# Field and experimental data together with computational models reveal how cleaner fish adjust decisions in a biological market

**DOI:** 10.1101/2022.03.10.483727

**Authors:** Andrés E. Quiñones, Zegni Triki, Redouan Bshary

## Abstract

While it is generally straightforward to quantify individual performance in cognitive experiments, identifying the underlying cognitive processes remains a major challenge. Often, different mechanistic underpinnings yield similar performances, and Lloyd Morgan’s cannon warrants acceptance of the simpler explanation. Alternatively, when the different mechanisms interact with environmental conditions, variation in performance across environments might allow to statistically infer the mechanism responsible. We illustrate this point by fitting computational models to experimental data on performance by wild-caught cleaner fish *Labroides dimidiatus* in an ephemeral reward task, as well as cleaner and client fish densities from the locations of capture. Using Bayesian statistics to fit the model parameters to performance data revealed that cleaner fish most likely estimate future consequences of an action, while it appears unlikely that the removal of the ephemeral reward acts as psychological punishment (negative reinforcement). Incorporating future consequences also yields performances that can be considered the result of locally optimal decision-rules, in contrast to the negative reinforcement mechanism. We argue that the combination of computational models with data is a powerful tool to infer the mechanistic underpinnings of cognitive performance.

**Author summary:** Performance in behavioural experiments is often used to assess the cognitive abilities of animals. However, animals can get to the same outcome in alternative ways. Thus, the outcome of the experiments does not tell us how animals achieve their performance. In order to overcome this limitation, we used a set of computational models, which provide predictions on how alternative cognitive mechanisms perform in the face of varying environmental conditions, and compare their predictions to performance data from individuals that experienced different conditions throughout their life. Our study system is the cleaner fish *Labroides dimidiatus*, a coral reef fish that feeds off ectoparasites and dead tissue from the skin of other reef fishes. Cleaner fish often have to choose among alternative clients seeking their cleaning service, and the optimal choice is often hard to achieve. Our performance experiments mimic the natural choices the cleaner fish make. The combination of experiments and computational models points to cleaner fish abilities to account for the future effect of their choices by estimating the long term expected value of those choices. Estimating long term value is a mechanism involved in human foresight.

## Introduction

Often alternative cognitive mechanisms yield similar behaviour and/or cognitive performances. This poses a problem for disentangling the mechanistic underpinnings of behaviour. This is particularly clear in research aimed at discovering between species variation in *higher* cognitive abilities; or in other words, research on whether non-human animals show cognitive abilities believed to be uniquely human. For instance, when researchers try to find *mental time travel-like* behaviour, they usually come up with clever experiments to show the behaviour displayed requires inferences made through past events[1]. However, they often face the challenge of alternative scenarios where simpler explanations, like classic associative learning, can bring about the observed behavioural outcome [2]. Similarly, attempts to demonstrate the presence of *theory of mind* in non-human animals face objections justified by alternative mechanisms underpinning similar behavioural results [3]. Such controversies are usually settled by using the principle of parsimony and its cognitive version, Lloyd Morgan’s cannon, which states that the simpler explanation (mechanism) should be accepted. Ideally, however, alternative hypotheses should be evaluated in light of their explanatory power. One potential dimension that could allow the variation necessary among distinct cognitive mechanisms is environmental variation. Controlled laboratory conditions often mask between and within species variation that arises among different environments. Thus, if predictions from the alternative mechanisms depend on environmental conditions, it is possible to assess their relative explanatory power by performing cognitive test in the context of different environmental conditions.

From an evolutionary perspective, mechanisms are likely selected because of how they allow individuals to respond to environmental variation. For example, biological market theory predicts that the exchange rate of goods and/or services traded between cooperative partners adjusts to the law of supply and demand, when individuals have some degree of partner choice [4]. Supply and demand conditions, which typically depend on the abundance of the species involved, certainly vary in time and space. Therefore, natural selection should favour the ability to flexibly adjust decisions and behavioural output to current market conditions. Indeed, such adjustments have been repeatedly documented [5]. In animals, an obvious general candidate mechanism for the strategic adjustment is the cognitive machinery. However, it is not clear which cognitive mechanisms allow individuals to adjust their behaviour to the varying conditions. More specifically the question arises to what extent mechanisms beyond basic associative learning may be involved.

One example of strategic adjustment in a biological market is the marine cleaning mutualism involving the cleaner fish *Labroides dimidiatus* and ‘client’ fish. Client fish seek cleaner fish services at their territory (so-called “cleaning station”) and offer themselves as food patches to get their ectoparasites removed, which provides cleaners with food and clients with improved health [6–9]. Given the capacity of some client fish to swim larger distances and access multiple cleaning stations while others access the only cleaning station in their territory, it is crucial to categorize clients as either “visitors” or “residents,” respectively. During cleaning interactions, a cleaner fish often face a choice between a visitor and a resident client seeking its cleaning services simultaneously. Visitors have the option to switch to another cleaner fish if being made to wait, while residents must wait for inspection. Indeed, visitors have been observed to use their partner choice option in that way [10], which may explain why cleaners give visitors service priority in a field study in the Red Sea [11].

When researchers aimed at testing wild-caught cleaner fish, as well as individuals from other species, abilities to prioritize visitor over resident clients in a lab-based paradigm, they used Plexiglas plates of different colours and shapes offering the same amount of food as surrogates for visitor and resident clients. One plate is made to behave like a resident, i.e. it remains until the cleaner fish has eaten off it. The other plate was made to behave like a visitor, i.e. it is removed if not inspected first. This laboratory paradigm originally inspired by the cleaning market of cleaner fish is more generally referred as the ephemeral reward task. Individuals are assessed in their capacity to develop a preference for the ephemeral option, which gives them access to two reward units. Cleaners learned to prefer the visitor plate and hence obtained the double amount of food [12]. Furthermore, adult cleaner fish outperformed various primates as well as rats and pigeons in this original version of the ephemeral reward task [13,14]; yet the African grey parrots solve this task as well [15]. This between variation in performance, and primates’ known cognitive abilities [16], shows that performance in the task cannot be readily explained by classifying species as having high and low cognitive abilities. Instead, this between species variation suggests that solving the ephemeral reward task requires a set of cognitive mechanisms that are not present in all the tested taxa, or were not used under the experimental set-up.

Recent research on the cognitive abilities needed to solve the ephemeral reward task has used a broad approach as proposed by Shettleworth [17], who defined cognition as including all ways in which animals acquire information through the senses, process, retain and decide to act on it. Indeed, perception of relevant cues is of major importance for individual performance. For example, some animals find relevant information in the plates [18] while others find it in the food [19,20]. However, identifying cues as salient is not the only challenge of the ephemeral reward task, as revealed by proximate learning models. Such models allow varying the cognitive tool kit and evaluating which minimal kit is necessary to solve the task at hand (e.g. [21]). Applied to the ephemeral reward task, learning models showed that basic reinforcement learning does not suffice to solve the ephemeral reward task [22,23]. This is particularly so when models assume the more complex natural situation in which cleaner fish face not only resident-visitor pairs but also visitor-visitor and resident-resident pairs, as well a resident alone or a visitor alone. To be able to give visitors priority over residents, cleaners need to be able to assess a client’s value separately for the three possible scenarios (alone, paired with a fish with the same strategic option, paired with a fish with the alternative strategic option) [23]. The ability to distinguish and value one stimulus differently alone from compound versions of it has been termed configurational learning, chunking, or segmentation (see references in [22]).

In addition to configurational learning, cleaners also need to account for the future consequences of current decisions. In the model by Quiñones *et al.* [23], this could be achieved in two non-mutually exclusive ways: through low temporal discounting of future effects, also termed ‘chaining’ [24]; and/or through perceiving a visitor client leaving as psychological punishment (i.e. as a negative reinforcer). Low temporal discounting is when individuals include in their valuation of an action the reward effects that this will have in the future. This is done by combining in a single valuation the reward obtained in the current time with all the reward that comes after, discounting for how far in the future reward is accrued. ‘Chaining’ the reward of these different time steps allows individuals to take actions that increase the long-term reward at the sacrifice of short term considerations [24]. Even though, ‘chaining’ can be readily implemented computationally in learning models [24,25], cognitively it seems to be a complex adaptation [2]. On the other hand, using client behaviour as a negative reinforcer is, in principle, easier to implement. Negative reinforcement is part of the ubiquitous learning mechanism termed operant conditioning [26,27]. Thus, the standard logic of Lloyd Morgan’s cannon demands that operant conditioning as the simpler explanation is to be accepted by default. Ideally, however, the two mechanisms should be evaluated in light of how well they explain the available data. Note that different fields interested in cognition and decision making use different words to refer to negative reinforcers [23,25]. Here, for the sake of simplicity and clarity, we will use the word ‘penalty’ to refer to this mechanism which includes a negative reinforcer.

Over the last decade, our team tested over a hundred wild-caught cleaner fish in the exact same paradigm of the ephemeral reward task [13,28–31]. These fish often come from different reef locations around the study field site at Lizard Island, Great Barrier Reef in Australia. Further investigation of the local eco-sociological conditions revealed that cleaner and client fish population densities have a substantial impact on cleaner fish performance in the task. Cleaner fish from reef sites with relatively low densities were more likely to fail at solving the task [28–30]. The explanation for this pattern is twofold. First, low cleaner fish population densities imply low supply and high demand for cleaning services. Under such conditions, there are fewer occasions under which failing to give priority to a visitor translates to an empty cleaning station. Second and also related to the state of the cleaning market, visitor clients at low cleaner fish population densities are less likely to exert their partner choice options and are hence more likely to wait for inspection if not given cleaning priority [31]. In the absence of partner choice cleaner fish should not give priority to visitors any more.

The interplay between the underlying cognitive machinery and local ecological conditions apparently generate the documented variable performance among individuals of the same species. Our approach of fitting the computational model to the empirical data on fish densities and cleaner fish censuses and performance in the ephemeral reward task aimed at: (i), determining which mechanism cleaner fish use to incorporate future consequences of current decisions by testing whether chaining, penalty, or a combination of both best explains their performance; (ii) determining whether the two mechanisms differed with respect to the ecological conditions that are likely to cause high versus low performance in the ephemeral reward task. Additionally, we assessed which mechanism yields optimal performance patterns. Relying on the logic of biological market theory, we predicted that appropriate performance is to show a low preference for visitors under low local cleaner-to-client ratio.

## Methods

### The model

The model consists of a set of individual-based simulations where individuals, representing cleaner fish, face a series of choices between two options, which simulate the natural conditions of the cleaning market. Individuals experience a series of discrete time points in which they face different ‘states,’ defined by the number and category of client fish (visitor or residents) inviting for cleaning services. There are six possible states: zero clients, one resident, one visitor, resident-resident, visitor-visitor, and resident-visitor. The probability of each state is largely determined by the relative abundance of cleaner fish, residents and visitors, but to some degree by cleaner fish choices when it faces the resident-visitor combination. This is because residents are willing to queue for cleaning service; while visitors leave the queue (with a certain probability) when made to wait. Individuals obtain a fixed reward from cleaning a client fish regardless of the category. Every time individuals face and make a choice they update the probability of making that same choice. The update is based on the difference between the expected reward and the obtained reward (prediction error) [25,32]. Given the new information, the update is carried in the direction that leads to more reward being obtained, given the new information. In the long run, the probability of choosing a visitor over a resident converges in the model. To which probability the model will converge depends on the relative abundance of cleaners, visitors and residents; as well as on the probability of visitors leaving the cleaning station when made to wait. Further details of the model implementation can be found in Quiñones *et al* [23].

The model shows that agents need to find a way to incorporate future consequences of current choices. In the model, this could be achieved with either of two parameters that could also work together. First, *γ* measures how much individuals include future rewards in their decision updates. If *γ* = 0, individuals only use the immediate reward obtained from a cleaning interaction. As *γ* increases, individuals include more the reward obtained from the subsequent choices. That amounts to calculating and using for decision making the future expected rewards of an action (chaining). Second, *η* measures how much individuals include in their reward the fleeing behaviour of visitors as a negative component (penalty). Both of these parameters allow individuals to use in their estimates the future effects of their choices.

### Empirical data

The empirical data were collected between 2010 and 2019 always during the austral winter months June to August from a total of five study reef sites (Corner Beach-CB, Horseshoe-HS, Mermaid Cove-MC, Northern Horseshoe-NHS, and The Crest-TC) at Lizard Island (14.6682°S, 145.4604°E), Great Barrier Reef, Australia. The data consist of three sets: fish censuses, field observations of cleaner-client interactions to quantify the probability of visitors leaving if made to wait, and the performance of wild-caught cleaner fish in the ephemeral reward test. In total, we have twelve site/year data sets for fish censuses and corresponding performance in the lab test. Thus, some sites were sampled more than once. To estimate the population density of cleaner fish and their clients at a given site in a given year, Triki *et al.* [30] used a series of ten transects of 30*m* each. Observers swam along the transect lines placed on the reef and first counted the visible large-bodied adult fish (species with total length *TL* ≥ 10*cm*) including cleaner fish on a width of 5*m*, and then on the return individuals of small-bodied fish species (*TL* < 10*cm*) on a width of 1*m* (see Triki et al. 2018 for further details on fish censuses data collection). We then scaled the counts of cleaner fish, small-bodied, and large-bodied clients fish densities per 100*m*^2^. From the study by Triki *et al* [30] cleaner fish and large-bodied client populations densities were highly correlated, and only the former was hence used in the analyses as representative of both measures.

The field observation data consisted of video recordings/encodings of the cleaner-client cleaning interactions. There were videos from eight cleaners per site/year of a duration of 30*min* each. Triki *et al* extracted information from every event wherein a visitor client was made to wait in favour of another client (visitor or resident), and noted whether or not the visitor left or queued for the cleaning service [30,31].

The cognitive performance data was from a total of 120 cleaners (10 individuals per 12 site/year) tested in the ephemeral reward task [30,31]. Authors housed all captured cleaners individually in glass aquaria ( 62*cm*×27*cm*cm×37*cm* ) and provided them with PVC pipes (10*cm*cm×1*cm*) as shelters. The task consisted of exposing the cleaner fish to substitute models of client fish in the form of two *Plexiglas* plates offering the same amount of food (one item of mashed prawn). The two plates differed in colour and pattern (horizontal green stripes or vertical pink stripes) but had equal size (10*cm*cm×7*cm*). Importantly, the two plates played different roles as either a visitor (ephemeral food source) or resident (permanent food source). That is, if a cleaner fish inspected the resident plate first, the experimenter withdrew the visitor plate out of the aquarium as a consequence. Choosing first the visitor plate, however, granted access to both plates. The equal size of the plates forced cleaner fish to learn to give service priority to the visitor plate based solely on the behaviour-cue of the plates rather than size-cue [18]. Triki *et al.* [30,31] tested the fish for a maximum of 200 trials with 20 trials a day, 10 trials in the morning and 10 trials in the afternoon. They randomized and counterbalanced the plates’ spatial location (i.e. left or right) between trials. Similarly, they counterbalanced the plates’ decoration (colour and pattern) and the plates’ role (visitor or resident) between the tested fish. In the original studies, once a fish reach a learning criterion, that is, performing significantly above chance level (> 50, *p − value* ≥ 0.05), they passed to a reversal version of the task where the roles of the visitor/resident Plexiglas plates were swapped (see Triki et al. 2019, 2020). Here, we used instead a subset of these data in order to have an idea of cleaner fish preferences for the visitor plate, even if they do not reach the learning criterion within 200 trials. To do so, we first extracted the trial-by-trial outcomes from the last two sessions (20 trials) of those who never reached the learning criterion for visitor plate (*N* = 45 cleaner fish). For those who reached the learning criterion at some point during the test and passed to a reversal phase, we extracted the trial-by-trial outcomes from the last session (10 trials) before passing to reversal and the last 10 trials they were exposed to in the test (*N* = 75 cleaner fish). We chose a combination of initial and reversal to quantify preference for the visitor client, given that it matches well the criteria chosen in previous analysis of the ephemeral reward task experimental set-up [30,31]. In the supplementary material (Fig. S1) we show how our choice maps to the previously used criteria. For comparison, we also show a quantification of visitor preference based only on the initial phase of the task.

### Ethics statement

The Animal Ethics Committee of the Queensland government (DAFF) approved the project (CA 2016/05/970 and CA 2017/05/1063)

### Statistical analysis

The aim of the analyses is to fit the key model parameters *γ* and *η*, to the empirical data from Triki *et al.* [30,31] to test whether each or a combination of these effects is a better explanation for the pattern seen in the data. We used the ecological variables: cleaners, visitor clients, resident clients abundances and visitor clients leaving probability, as input to the models. As the response variable, we used cleaners’ preference for visitors over residents in the ephemeral reward task. Finally, we used the preference for visitors resulting from the model simulations as the prediction for the response variable. We kept all other parameter values used for the model simulations constant, see Table S1.

To capture with the model the relationship between the ecological variables and cleaner fish preferences for visitors, we needed to scale the absolute population densities of cleaner fish from the empirical data to a measure of relative abundances that captures client visitation patterns. This is because, in the model, relative abundances of clients define not only the probability of residents and visitors but also how often the cleaning station is empty (e.g. there are no clients to be cleaned). The frequency with which clients visit the station is another variable, which in nature may vary among different client species depending on their ectoparasite loads. We do not have field estimates for species-specific parasite loads, especially not as a function of the site. In order to control for these aspects, we computed a measure of relative cleaner fish abundance for each reef site relative to absolute abundances and multiplied it by a scaling constant that changes the range of the variable. This scaling constant is hence meant to capture variation in the market conditions driven partly by cleaner fish abundance. We fitted the value of the scaling constant as part of the statistical inference. As for the visitor and resident abundances, we computed a relative measure with respect to the total client abundance, and weighted that by the rescaled relative absence of cleaners (1 − relative cleaner abundance). Thus, all three measures of relative abundance sum up to 1, and can be used in the model as a proxy for the probability of having different options in the cleaning station. Note that we introduced the scaling constant to control for variation that is not captured by the model; its parameter distribution does not offer biological insights.

Once we calculated the relative abundances, we obtained predictions from the model for each one of the locations and ran the Markov Chain. We started the chain with random values for the parameters of interest, then ran the computational model once for every reef site using as input the ecological explanatory variables. The model outputs the probability of choosing the visitor for each location *p_i_*, where *i* is the index for the 12 locations. Assuming a binomial distribution, the probability that each of the cleaner fish 20 choices in the ephemeral reward task was generated by the model is given by 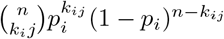; where *k_ij_* is the number of times that cleaner fish *j* in reef site *i* chose the visitor over the resident; and *n* = 20 (due to our choice of using 20 choices per cleaner fish). By taking the natural logarithm and summing over all the cleaner fish and reef sites, we obtained the log-likelihood of the data given the model and parameters. We then proposed a new set of parameters drawn from a uniform distribution centred around the old parameter set. The amplitude of the uniform distribution used for each parameter can be found in Table S1. Subsequently, we ran the model and calculated the likelihood with the new parameter set. We then used the ratio of the two likelihoods to choose which parameter set to keep. New parameter sets with a higher likelihood than the old set replaced old ones, and those with a lower likelihood replaced current ones with a probability equal to the log-likelihood ratio. Given that we only used the likelihoods in the decision, we used an uninformative prior. Once we decided whether the new parameter set would replace the old one, we ran the model again to sample the likelihood distribution of the parameter set. We then started the cycle again by proposing a new set of parameters and repeated the process for 1*e*^5^ steps. We ran 5 independent chains, discarded the first 1000 samples of each chain as burn in, and after that, we kept 1 in every 100 samples to avoid autocorrelation. The collection of parameters sets kept in all chains approximates the posterior distribution. We coded the model as well as the fitting algorithm in *c++*, the diagnostics and visualization in R [33]. All codes are accessible at <DOI: 10.5281/zenodo.6338073>

To compare the fit of the three alternative models, we used the distribution of pseudo *R*^2^ proposed by Mcfadden [34]. Mcfadden’s pseudo *R*^2^ is a standard measure of fit for logistic regression. In that context, *pseudo* − *R*^2^ uses the log-likelihood of the data given the model, relative to the log-likelihood of the data given a model without covariates, as a measure of fit. Our model is not a logistic regression, therefore we measured the *pseudo* − *R*^2^ in relation to the log-likelihood of a model with parameters *γ* and *η* set to zero. This, in practice, amounts to a model that has a neutral preference between the two options. We computed the pseudo *R*^2^ for all the samples of the posterior from the MCMC. Thus, we used these distributions of pseudo *R*^2^’s as a measure of fit.

## Results

Estimation of parameter values for the three models (full model, chaining and penalty), support chaining as the only mechanism cleaners use to account for the future effects of their actions; and thus to solve the ephemeral reward task. In the estimation of the parameter values of the full model, which includes both chaining and penalty, the bulk of the marginal posterior distribution of *η* which controls the strength of penalty is around 0 (Fig. 1). As for *γ*, controlling chaining, the 95% confidence intervals also includes zero, but the mode of the posterior is around 0.5 (Fig. 1, a). In the chaining model, where *η* is set to zero, the distribution of *γ* shifted to higher values, zero is no longer part of the 95% credible interval of the parameter (Fig. 1, c). In contrast, when we look at the model with only penalty, the posterior distribution of *η* is still centred around zero (Fig. 1, d). Thus, the analysis of the estimates of individual parameter values in the three models only supports a strong effect of chaining. Furthermore, the comparison of the models’ fit favours the chaining model. In panel e of figure 1 we show the distribution of *pseudo* − *R*^2^ calculated using samples from the posterior distributions shown before. Note, *pseudo* − *R*^2^ can have negative values, which is when the log-likelihood of the model is lower than that of a model that triggers neutral preferences. Even though the peak of the three *pseudo* − *R*^2^ distributions were not very different, the model with only chaining produced a distribution of *pseudo* − *R*^2^ where more values were positive (to the right of the black line in Fig. 1 f). This shows that accounting for variation in the parameter estimates the model with chaining gives a better fit to the data, despite having one parameter less than the full model. We have not shown here the marginal posterior distributions of the scaling constant, given that they do not bring biological insight. Their visualizations can be found in the supplementary material (Fig. S2), as well as the diagnostics of the MCMCs (Figs. S3,S4,S5).

**Fig 1.**
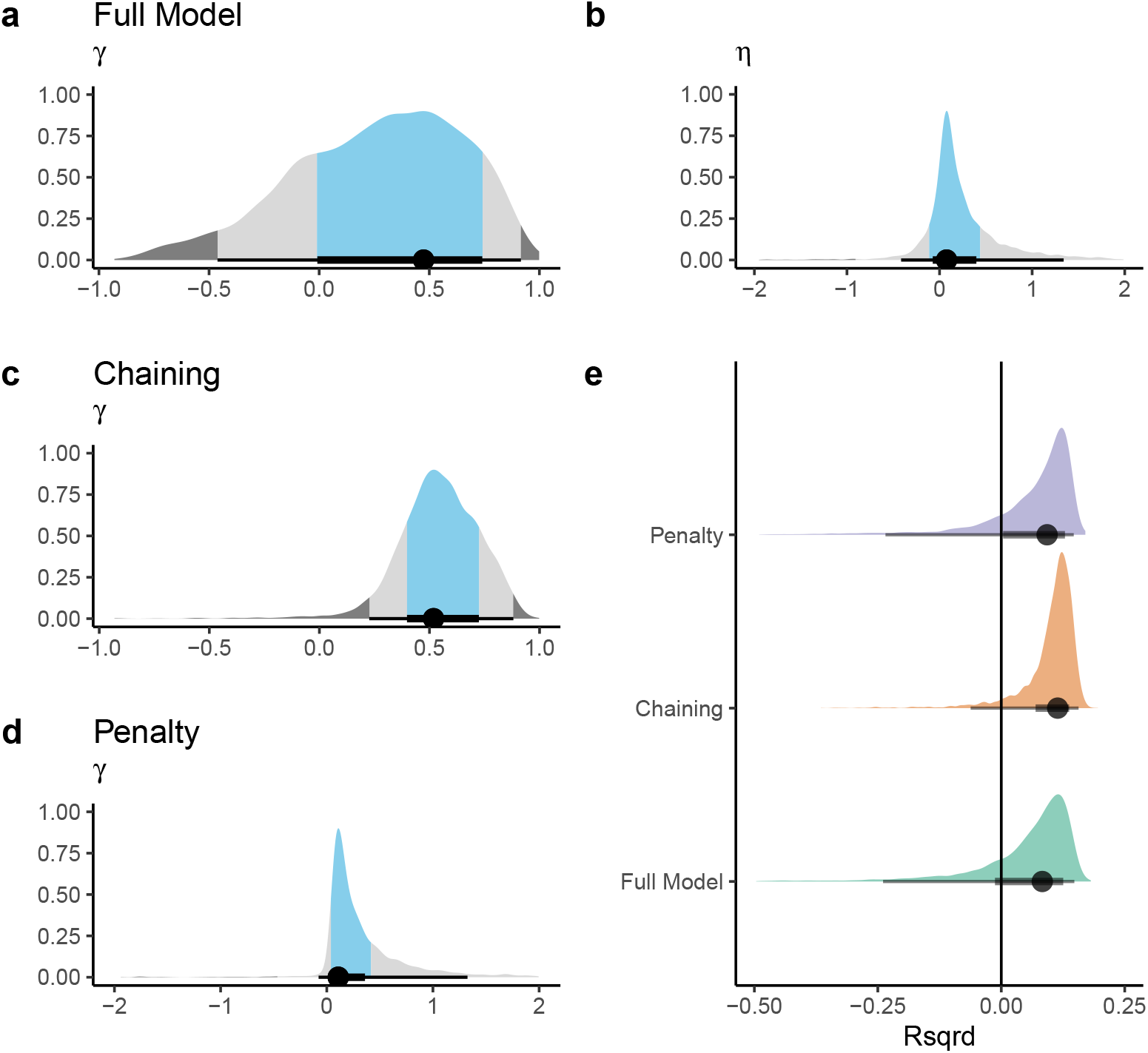
Posterior distributions for parameter values *γ* and *η* for the three models (Full model, chaining only and penalty only). We show the kernel density estimates, below the mode (black dot) and the 65 (light blue shade) and 95% (grey shade) highest posterior density interval for the two parameters. On the top, panels a and b show posterior distributions for a full model, including chaining (*γ*) and penalty (*η*). Panel c, shows the *γ* estimate from model with only chaining. Panel d shows the *η* estimate from a model with only penalty. Panel e, shows a measure of fit for all models, namely the distribution of pseudo-*R*^2^ obtained from sampling the posterior distribution of parameter values.

The main reason for chaining and penalty to give different predictions is the way that cleaner fish relative abundance influences the preference for the visitor clients. Visitor leaving probability has a similar positive effect on the probability of choosing the visitor clients on all three models, they all predict an increase preference for visitors as the visitor probability increases (Fig. 2). In contrast, cleaner fish relative abundance has a different effect in the model with only chaining, compared to the other two models, full model and penalty. In the model with only chaining, only intermediate cleaner fish abundance triggers a preference for the visitor clients (Fig. 2 c). In both models with penalty, both intermediate and low cleaner abundance triggered a preference for the visitor (Fig. 2 a,e). Note, however, we calculated preferences shown in figure 2 left panels by using only the mode of the posterior distributions, and by holding constant the balance between resident and visitors’ abundances. Panels on the right, show how close predictions are from the observed data, allowing the balance between client types to vary and using a set of samples from the posterior distribution.

**Fig 2.**
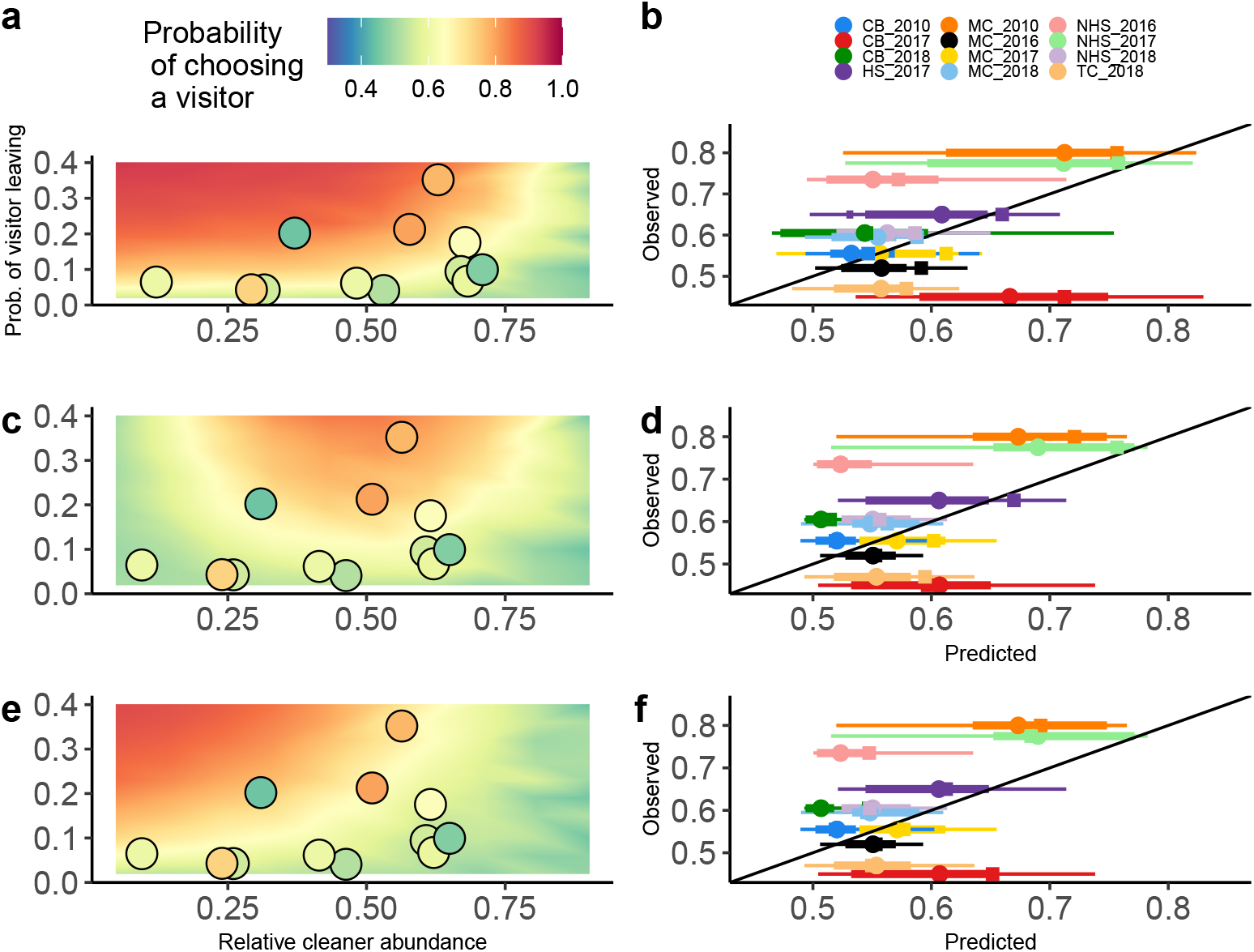
Observed and predicted probability of choosing a visitor. Left-hand side panel: colour contour shows the prediction from the learning model using the mode of the posterior distributions of parameters recovered by the statistical analysis. Dots show the frequency of visitor choices for the 12 reef sites, as well as the corresponding relative cleaner fish abundance (x axis) and frequency of visitors clients leaving the cleaning station (y axis). Right-hand side panels: Variation of the predicted probabilities of choosing a visitor over a resident and their observed values for 12 locations. Circles show the mean prediction for each location from 100 samples taken from the posterior distribution. Thick and thin bars show the 66 and 95% credible interval, respectively, taken from those posterior samples. Squares show the predictions used for the panel on the right-hand side. Colour coding denotes different reef site/year of the data collection (see Empirical data section). The black line corresponds to a perfect match between observed and predicted probabilities. Upper panels (a and b) show predictions from a model including chaining and penalty; middle panels (c and d) from a model with only chaining; lower panels (e and f) from a model with penalty only.

## Discussion

In this study, our main aim was to unravel which of two potential cognitive mechanisms, chaining of events, penalty, or their combination, best explains wild-caught cleaner fish performance in the ephemeral reward task, while accounting for their ecological conditions. To evaluate the merits of each of these two mechanisms separately and combined, we considered cleaner fish performance in the lab test to have its origin from the rule these fish applied in their natural environment. That is, individuals that solved the task already had a preference for visitors clients and generalized this rule to the lab conditions once being familiar with the task.

While all three models captured well the positive relationship between visitor leaving behaviour and cleaner fish performance in the market task [30], only the chaining mechanism predicted that cleaner fish performance in the task should be low in habitats with low cleaner-to-client ratios, regardless of the visitor leaving probability. In contrast, models including negative reward predicted the highest performance in the ephemeral reward task when relative cleaner fish abundance is low, particularly together with a high probability of visitor leaving (Fig. 2). Low relative cleaner fish abundances mean the market has an excess of demand for cleaning services. In the models, this translates to a cleaning station that is frequently full. Thus, when a visitor leaves, it is likely that the cleaner fish will have access to another client in the next step. Therefore, there will not be much difference in future reward between choosing a visitor and a resident, and cleaners will not develop a preference for the visitor in these conditions. On the other hand, the effect of negative reward on cleaner fish preference is the opposite, as in a busy cleaning station, to that of chaining. cleaner fish will get more often the resident-visitor state and will develop a preference for the visitor faster. At high cleaner fish abundances, the resident-visitor state becomes so rare that neither mechanism is very efficient at generating a preference for visitors. When facing the resident-visitor choice, it is still best to choose the visitor; however, the learning machinery will not be able to develop this preference efficiently. Overall, the models suggest that chaining is the cognitive mechanism that allows cleaner fish to adaptively adjust to their biological market ecological conditions.

Previous research showed that cleaner fish living at high population densities and giving service priority to the visitor plate in the ephemeral reward task, as well as cleaner fish living at low densities but denying service priority to the visitor plate possess larger forebrains; a key teleost brain region associated with behavioural flexibility and social intelligence. Those failing to show optimized decision-rules given their local ecological conditions had relatively smaller forebrains [31]. Triki *et al.* refer to the former as socially competent cleaner fish, while the second group as socially incompetent cleaner fish. Social competence is the ability to optimise social behaviour to the available social information [35–37]. Our analyses yielded no evidence that the difference in social competence with respect to the local ecological conditions and associated brain morphology, found by Triki *et al* [31], is due to the mechanism used to incorporate future consequences. It is conceivable that high performing individuals from low population densities reef sites use negative reinforcement instead of chaining, but in that case, negative reinforcement should have explained at least part of the data. Configurational learning or chunking [38,39], the second component necessary to solve the ephemeral reward task [23], was not varied in the models we analysed here. However, while chunking tendencies should vary to allow individuals to adapt to their local conditions [22,40], systematic differences in individual chunking tendencies would not explain how socially competent decisions vary as a function of relative abundance. Therefore, it remains currently unclear what cell-demanding mechanisms may cause variation in social competence that translates into site-specific variation in performance in the ephemeral reward task.

Our models are inspired by the general processes of associative learning where short term rewards are translated into decision making; thus, it ignores alternative channels of information that could be relevant in market-like situations. For example, the model does not investigate whether cleaner fish actually assess the frequency of client visits or a mean frequency of visitors leaving. The updating learning mechanism for the development of preferences works on a trial-by-trial basis. In the model, cleaner fish do not need to assess the actual state of the market, *i.e.* their abundance, the abundance of residents and visitors, and client visitation rate as an indicator of demand. They only need to assess the short-term consequences of their own decisions on food intake and chain them. Also, for the sake of simplicity, the model ignores the process by which cleaner fish discriminate residents and visitor clients. A model that accounts for this discrimination probably would involve the development of preferences for morphological or behavioural features that are statistically associated with visitors or residents. For example, visitors are on average larger than residents in body size [11], and contrary to residents, they are less likely to chase a cleaner fish that fails to cooperate and instead cheats its client by taking a bite of mucus [12]. Given these associations, chaining might produce the decision-rule “choose the larger client and/or the less aggressive client,” which is not a useful rule in the standard ephemeral reward task.

In conclusion, our study shows that variation in cognitive performance as a function of the local ecological conditions may set the stage for the use of mechanistic modelling to identify the cognitive processes underlying learning in animals. The combination overcomes the limitations of the general philosophy in animal cognition to apply the logic of Lloyd Morgen’s canon (Occam’s razor). Cognitive experiments with the aim of excluding basic reinforcement learning as a potential explanation (operant and/or classical conditioning) of performance often employ one trial experiments requiring animals to solve the task on the first possible occasion. For example, any theory of mind task needs to be solved in the first trial in order to exclude fast conditioning [3]. Similarly, subjects need to solve a social learning task on the first trial to accept imitation as a mechanism over stimulus/local enhancement. Such strict conditions are virtually never met. For example, potato washing by Japanese macaques, an iconic example of social learning, took several years to spread within the group [41], meaning that any learner had been repeatedly exposed to demonstrations before acquisition. Importantly, Galef [42] refuted imitation as a mechanism not simply because of the repeated exposure but because a (rather qualitative) analysis of the spread of potato washing across individuals did not follow the prediction based on imitation learning (see also [43]). In our case, the number of trials it took cleaners to learn the solution to the ephemeral rewards task would never allow excluding an important role of penalty based on the data alone. However, fitting model predictions to our comprehensive empirical data set revealed that a more complex mechanism, estimation of future reward, fits the data better.

## Supplementary material

**Fig S1.**
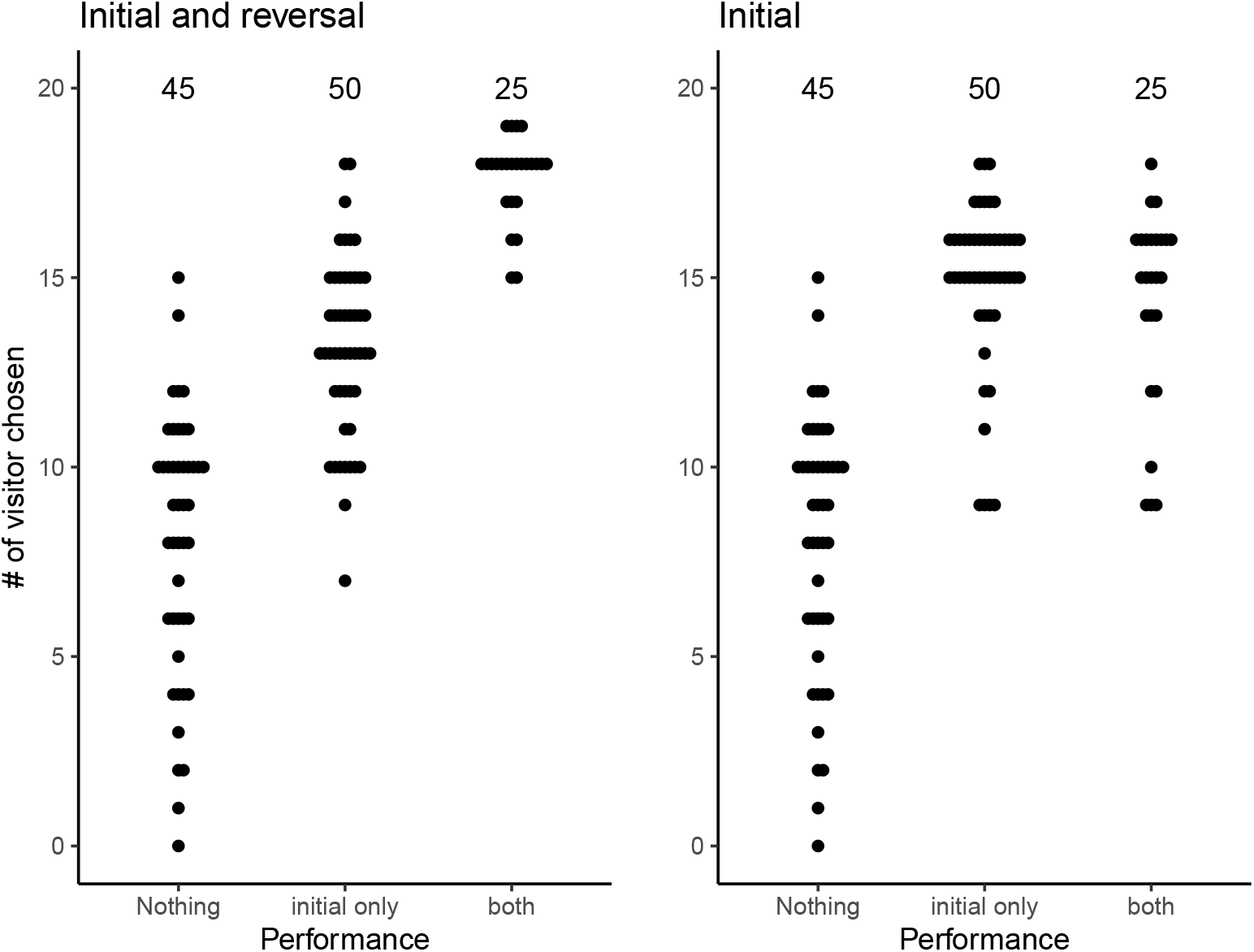
Relation between the response variable used in this study and the criteria used in previous studies to assess performance in the ephemeral reward task. In the x axis, we classified the performance of cleaner fish according to whether they developed a preference for the visitor in the initial round, in the initial and reversal, or none of them. In the y axis, we add the choices of two experimental sessions: panel on the left uses one session from the initial round and one from the reversal round when possible (as described in the main text); panel on the right uses two sessions from the initial round for all fish.

**Fig S2.**
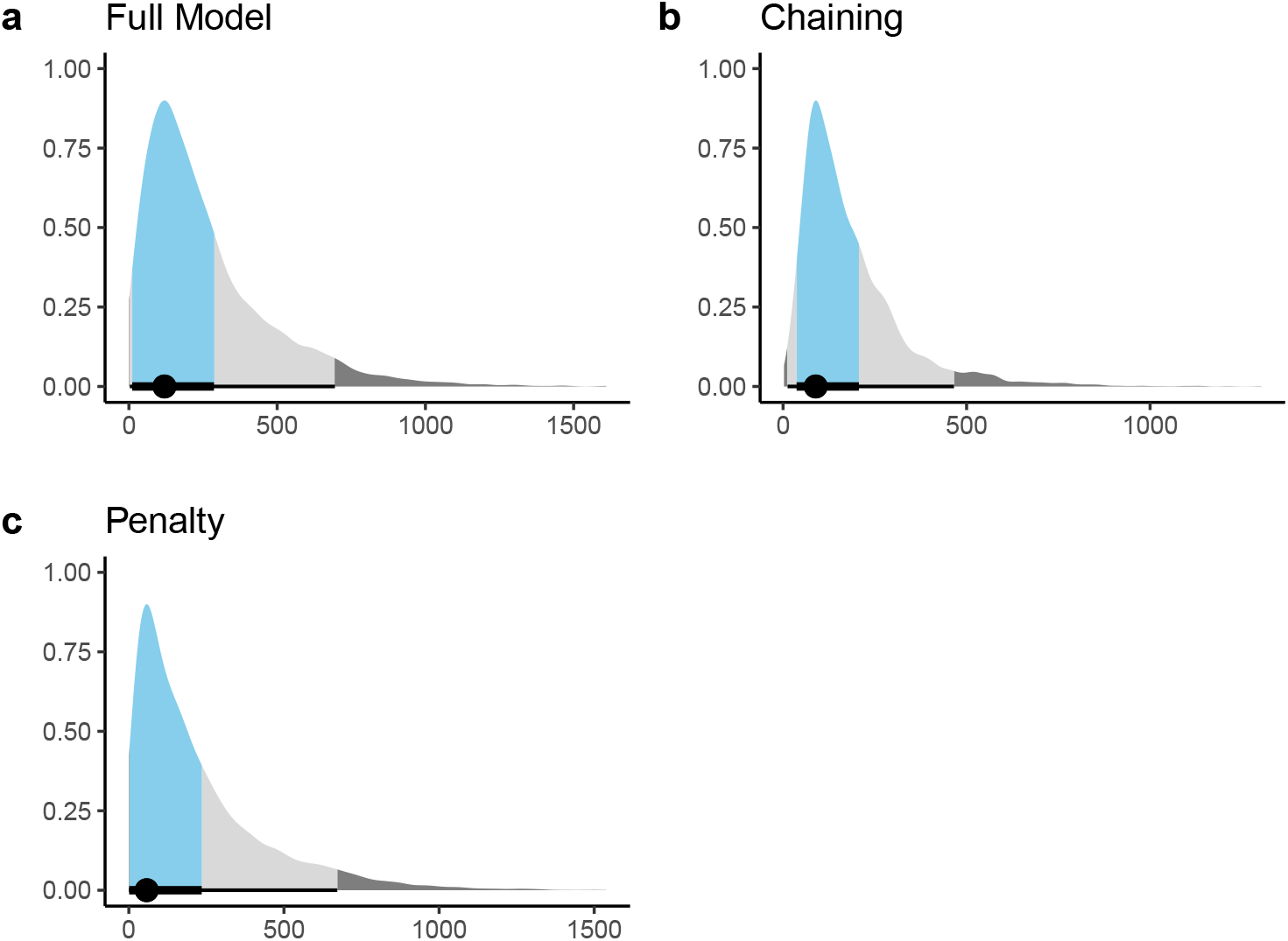
Posterior distributions for scaling constant for the three models (Full model, chaining only and penalty only). We show the kernel density estimates, below the mode (black dot) and the 65% (light blue shade) and 95% (grey shade) highest posterior density interval. On the top, panel a shows the posterior distribution from the full model; panel b from the model with only chaining; and panel c from a model with only penalty.

**Fig S3.**
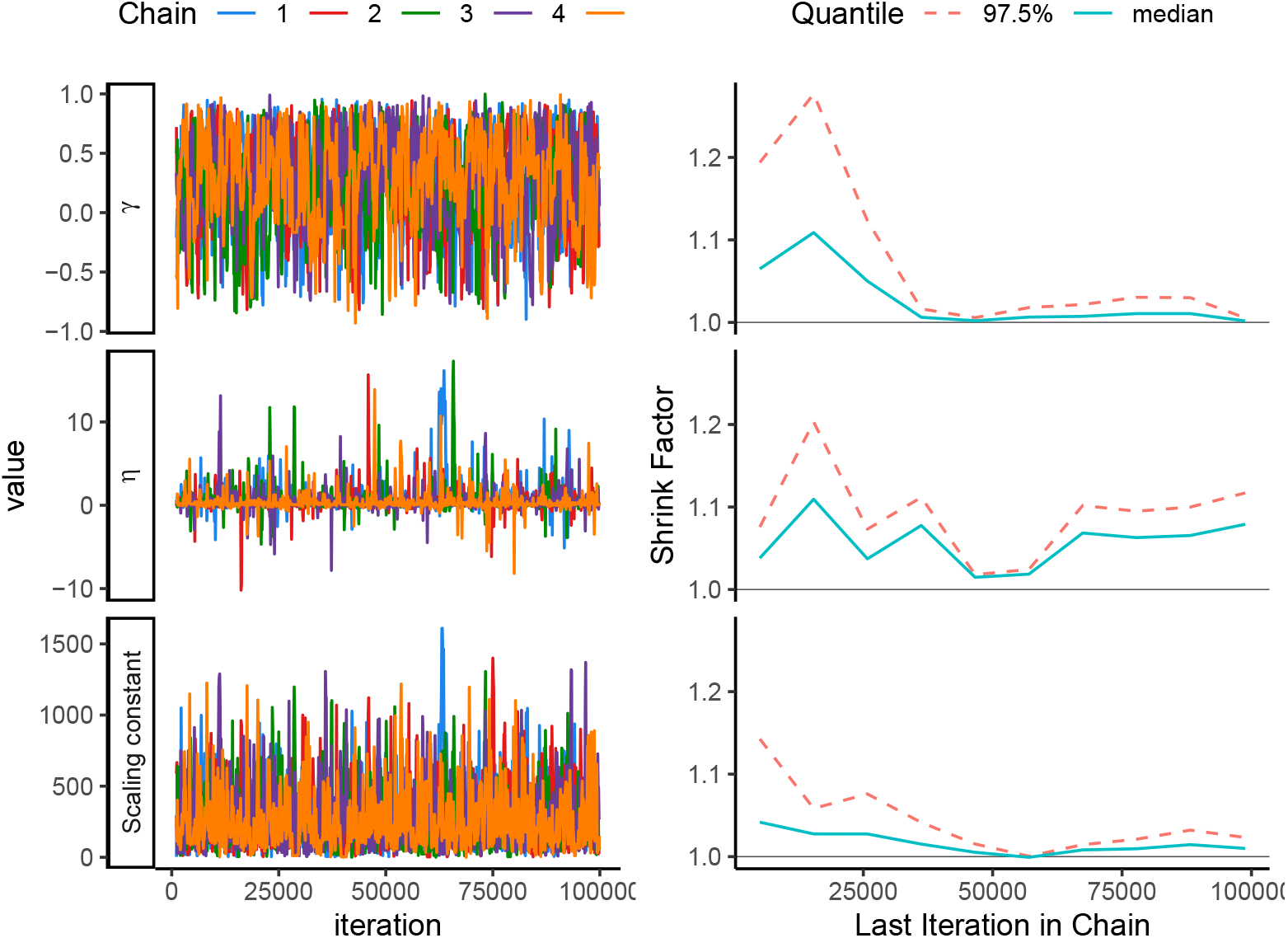
MCMC convergence diagnostics for the full model. On the left trace-plots, on the right changes along the chain of the Gelman and Rubin shrink factor.

**Fig S4.**
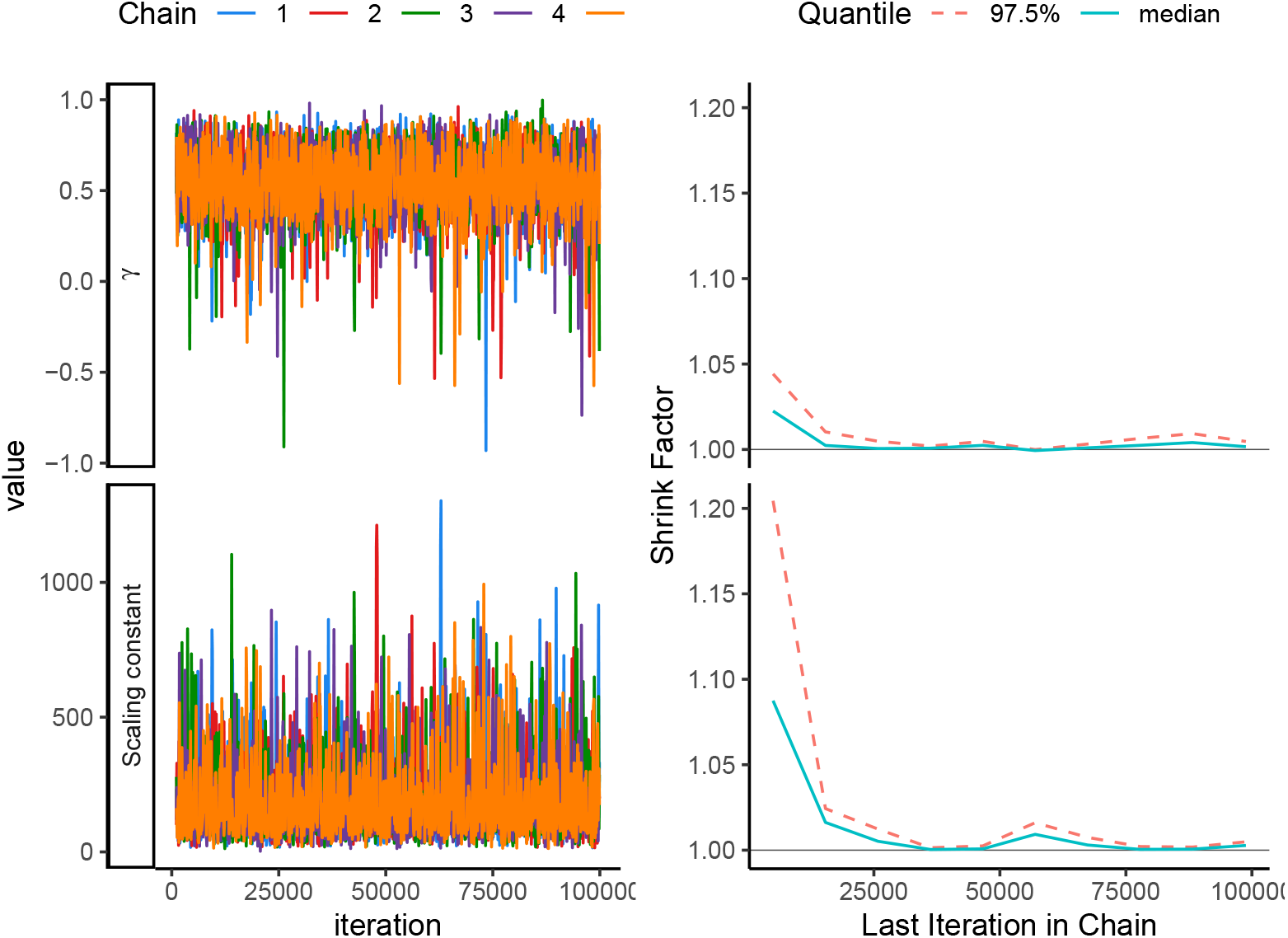
MCMC convergence diagnostics for the chaining model. On the left trace-plots, on the right changes along the chain of the Gelman and Rubin shrink factor

**Fig S5.**
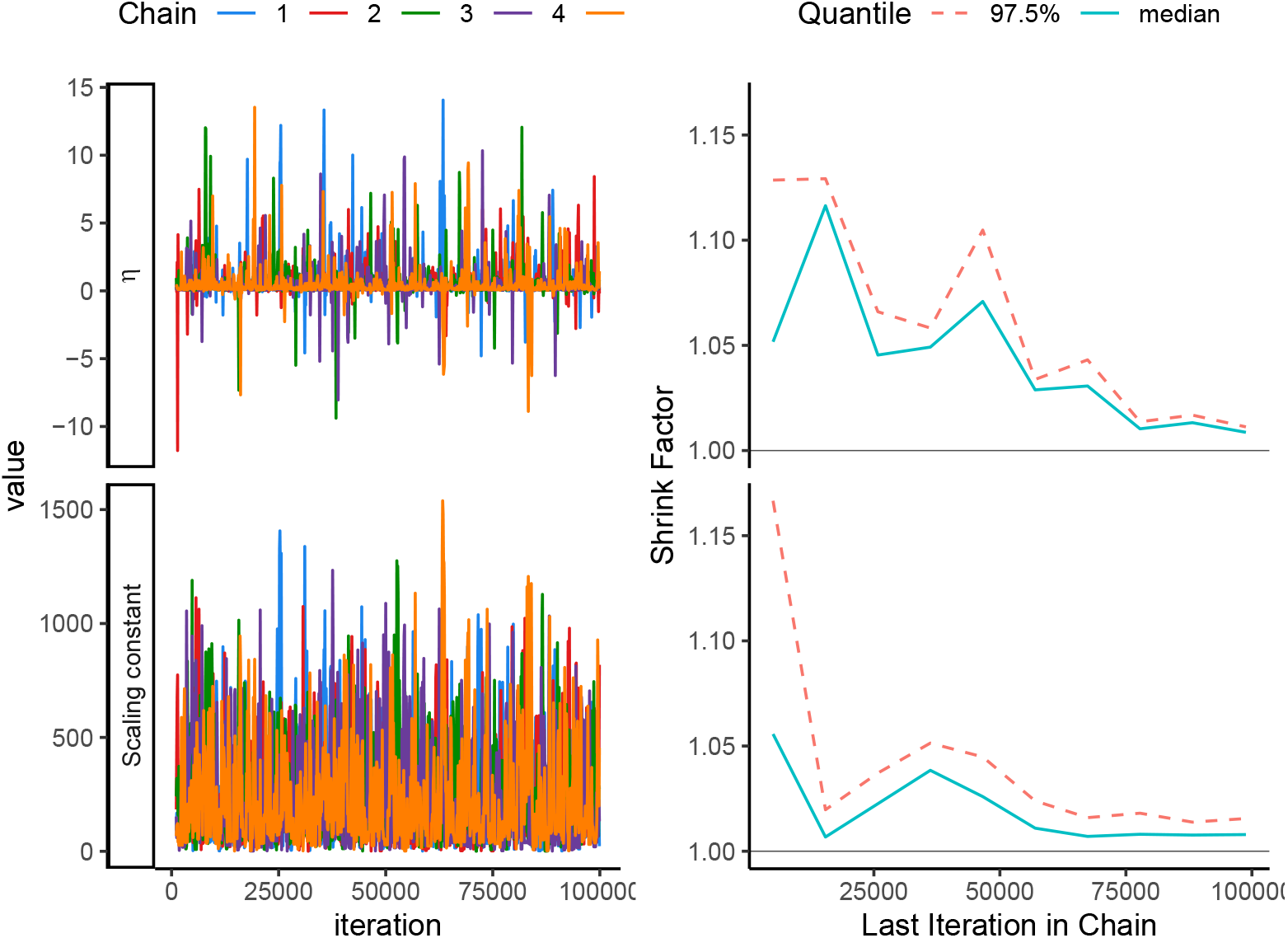
MCMC convergence diagnostics for the penalty model. On the left trace-plots, on the right changes along the chain of the Gelman and Rubin shrink factor

**Table S1.** Parameter values with which the model was run in the MCMC. *σ* refers to the amplitude of the perturbation kernel with the subscript indicating the associated parameter. New values were taken from a uniform distribution. *α* refers to the learning rate.

| Parameter | Value |
| --- | --- |
| Learning rounds | 10000 |
| Reward value | 1 |
| $\alpha$ | 0.05 |
| $\sigma_\gamma$ | 0.3 |
| $\sigma_\eta$ | 4 |
| $\sigma_{Sca.Const.}$ | 300 |
| Number of chains | 5 |
| Chain length | $1^5$ |

## References

1. Dally JM, Emery NJ, Clayton NS. Food-caching western scrub-jays keep track of who was watching when. Science (New York, NY). 2006;312: 1662–1665. doi:10.1126/science.1126539

2. Suddendorf T, Corballis MC. The evolution of foresight: What is mental time travel, and is it unique to humans? The Behavioral and Brain Sciences. 2007;30: 299–313; discussion 313-351. doi:10.1017/S0140525X07001975

3. Heyes CM. Theory of mind in nonhuman primates. Behavioral and Brain Sciences. 1998;21: 101–114. doi:10.1017/S0140525X98000703

4. Noë R, Hammerstein P. Biological markets. Trends in Ecology & Evolution. 1995;10: 336–339. doi:10.1016/S0169-5347(00)89123-5

5. Axén AH, Leimar O, Hoffman V. Signalling in a mutualistic interaction. Animal Behaviour. 1996;52: 321–333. doi:10.1006/anbe.1996.0178

6. Waldie PA, Blomberg SP, Cheney KL, Goldizen AW, Grutter AS. Long-Term Effects of the Cleaner Fish Labroides dimidiatus on Coral Reef Fish Communities. PLOS ONE. 2011;6: e21201. doi:10.1371/journal.pone.0021201

7. Ros AFH, Lusa J, Meyer M, Soares M, Oliveira RF, Brossard M, et al. Does access to the bluestreak cleaner wrasse Labroides dimidiatus affect indicators of stress and health in resident reef fishes in the Red Sea? Hormones and Behavior. 2011;59: 151–158. doi:10.1016/j.yhbeh.2010.11.006

8. Triki Z, Grutter AS, Bshary R, Ros AFH. Effects of short-term exposure to ectoparasites on fish cortisol and hematocrit levels. Marine Biology. 2016;163: 187. doi:10.1007/s00227-016-2959-y

9. Demairé C, Triki Z, Binning SA, Glauser G, Roche DG, Bshary R. Reduced access to cleaner fish negatively impacts the physiological state of two resident reef fishes. Marine Biology. 2020;167: 48. doi:10.1007/s00227-020-3658-2

10. Bshary R, Schäffer D. Choosy reef fish select cleaner fish that provide high-quality service. Animal Behaviour. 2002;63: 557–564. doi:10.1006/anbe.2001.1923

11. Bshary R. The cleaner fish market. In: Van Hooff JARAM, Hammerstein P, Noë R, editors. Economics in Nature: Social Dilemmas, Mate Choice and Biological Markets. Cambridge: Cambridge University Press; 2001. pp. 146–172. doi:10.1017/CBO9780511752421.010

12. Bshary R, Grutter AS. Asymmetric cheating opportunities and partner control in a cleaner fish mutualism. Animal Behaviour. 2002;63: 547–555. doi:10.1006/anbe.2001.1937

13. Salwiczek LH, Prétôt L, Demarta L, Proctor D, Essler J, Pinto AI, et al. Adult Cleaner Wrasse Outperform Capuchin Monkeys, Chimpanzees and Orang-utans in a Complex Foraging Task Derived from Cleaner – Client Reef Fish Cooperation. PLOS ONE. 2012;7: e49068. doi:10.1371/journal.pone.0049068

14. Zentall TR, Case JP, Berry JR. Early commitment facilitates optimal choice by pigeons. Psychonomic Bulletin & Review. 2017;24: 957–963. doi:10.3758/s13423-016-1173-8

15. Pepperberg IM, Hartsfield LA. Can grey parrots (Psittacus erithacus) succeed on a “complex” foraging task failed by nonhuman primates (Pan troglodytes, Pongo abelii, Sapajus apella) but solved by wrasse fish (Labroides dimidiatus)? Journal of Comparative Psychology (Washington, DC: 1983). 2014;128: 298–306. doi:10.1037/a0036205

16. Reader SM, Hager Y, Laland KN. The evolution of primate general and cultural intelligence. Philosophical Transactions of the Royal Society B: Biological Sciences. 2011;366: 1017–1027. doi:10.1098/rstb.2010.0342

17. Shettleworth SJ. Cognition, Evolution, and Behavior. 2 edition. Oxford ; New York: Oxford University Press; 2009.

18. Wismer S, Pinto AI, Triki Z, Grutter AS, Roche DG, Bshary R. Cue-based decision rules of cleaner fish in a biological market task. Animal Behaviour. 2019 [cited 30 Oct 2019]. doi:10.1016/j.anbehav.2019.09.013

19. Prétôt L, Mickelberg J, Carrigan J, Stoinski T, Bshary R, Brosnan SF. Comparative performance of orangutans (Pongo spp.), Gorillas (Gorilla gorilla gorilla), and drills (Mandrillus leucophaeus), in an ephemeral foraging task. American Journal of Primatology. 2021;83: e23212. doi:10.1002/ajp.23212

20. Prétot L, Bshary R, Brosnan SF. Comparing species decisions in a dichotomous choice task: Adjusting task parameters improves performance in monkeys. Animal Cognition. 2016;19: 819–834. doi:10.1007/s10071-016-0981-6

21. Dubois T, Pasquaretta C, Barron AB, Gautrais J, Lihoreau M. A model of resource partitioning between foraging bees based on learning. PLOS Computational Biology. 2021;17: e1009260. doi:10.1371/journal.pcbi.1009260

22. Prat Y, Bshary R, Lotem A. Modelling how cleaner fish approach an ephemeral reward task demonstrates a role for ecologically tuned chunking in the evolution of advanced cognition. PLOS Biology. 2022;20: e3001519. doi:10.1371/journal.pbio.3001519

23. Quiñones AE, Leimar O, Lotem A, Bshary R. Reinforcement Learning Theory Reveals the Cognitive Requirements for Solving the Cleaner Fish Market Task. The American Naturalist. 2019; 000–000. doi:10.1086/707519

24. Enquist M, Lind J, Ghirlanda S. The power of associative learning and the ontogeny of optimal behaviour. Royal Society Open Science. 2016;3: 160734. doi:10.1098/rsos.160734

25. Sutton RS, Barto AG. Reinforcement Learning: An Introduction. Introduction edition edition. Bach F, editor. Cambridge, MA: A Bradford Book; 2018.

26. Thorndike EL. Animal intelligence: An experimental study of the associative processes in animals. The Psychological Review: Monograph Supplements. 1898;2: i–109. doi:10.1037/h0092987

27. Skinner BF. The behavior of organisms;: An experimental analysis,. 1938.

28. Wismer S, Pinto AI, Vail AL, Grutter AS, Bshary R. Variation in Cleaner Wrasse Cooperation and Cognition: Influence of the Developmental Environment? Ethology. 2014;120: 519–531. doi:10.1111/eth.12223

29. Triki Z, Wismer S, Levorato E, Bshary R. A decrease in the abundance and strategic sophistication of cleaner fish after environmental perturbations. Global Change Biology. 2018;24: 481–489. doi:10.1111/gcb.13943

30. Triki Z, Wismer S, Rey O, Ann Binning S, Levorato E, Bshary R. Biological market effects predict cleaner fish strategic sophistication. Behavioral Ecology. 2019;30: 1548–1557. doi:10.1093/beheco/arz111

31. Triki Z, Emery Y, Teles MC, Oliveira RF, Bshary R. Brain morphology predicts social intelligence in wild cleaner fish. Nature Communications. 2020;11: 6423. doi:10.1038/s41467-020-20130-2

32. Rescorla RA, Wagner ARA. A theory of Pavlovian conditioning: Variations in the effectiveness of reinforcement and non-reinforcement. In: Black AH, Prokasy WF, editors. Classical conditioning II: Current research and theory. New York: Appleton-Century-Crofts; 1972.

33. R Core Team. R: A language and environment for statistical computing. Vienna, Austria; 2021. Available: https://www.R-project.org/

34. McFadden D. Conditional logit analysis of qualitative choice behavior. Frontiers in econometrics. 1974.

35. Taborsky B, Oliveira RF. Social competence: An evolutionary approach. Trends in Ecology & Evolution. 2012;27: 679–688. doi:10.1016/j.tree.2012.09.003

36. Bshary R, Oliveira RF. Cooperation in animals: Toward a game theory within the framework of social competence. Current Opinion in Behavioral Sciences. 2015;3: 31–37. doi:10.1016/j.cobeha.2015.01.008

37. Varela SAM, Teles MC, Oliveira RF. The correlated evolution of social competence and social cognition. Functional Ecology. 2020;34: 332–343. doi:10.1111/1365-2435.13416

38. Sutherland RJ, Rudy JW. Configural association theory: The role of the hippocampal formation in learning, memory, and amnesia. Psychobiology. 1989;17: 129–144. doi:10.3758/BF03337828

39. Miller GA. The magical number seven, plus or minus two: Some limits on our capacity for processing information. Psychological review. 1956;63: 81.

40. Kolodny O, Edelman S, Lotem A. The evolution of continuous learning of the structure of the environment. Journal of The Royal Society Interface. 2014;11: 20131091–20131091. doi:10.1098/rsif.2013.1091

41. Kawamura S. The process of sub-culture propagation among Japanese macaques. Primates. 1959;2: 43–60. doi:10.1007/BF01666110

42. Galef BG. The question of animal culture. Human Nature. 1992;3: 157–178. doi:10.1007/BF02692251

43. Hirata S, Watanabe K, Masao K. “Sweet-Potato Washing” Revisited. In: Matsuzawa T, editor. Primate Origins of Human Cognition and Behavior. Tokyo: Springer Japan; 2001. pp. 487–508. doi:10.1007/978-4-431-09423-4_24

